# INCLUSION CRITERIA UPDATE OF THE INTRALUMINAL ISCHEMIC MODEL IN RAT FOR PRECLINICAL STUDIES

**DOI:** 10.1101/114629

**Authors:** Héctor Fernández Susavila, Ramón Iglesias Rey, Antonio Dopico López, María Pérez Mato, Tomás Sobrino Moreira, José Castillo Sánchez, Francisco Campos Pérez

## Abstract

A proper occlusion of the medial cerebral artery (MCA) determined by laser Doppler during cerebral ischemia in rat models is an important inclusion criteria in experimental studies. However, a successful occlusion of the artery does not always guarantee a reproducible infarct volume which is critical to validate the efficacy of new protective drugs. In this study, we have compared the variability of infarct size in ischemic animals when the artery occlusion is monitored with laser Doppler alone and in combination with MRI during artery occlusion. Infarct volume determined at 24 hours was compared between animals with laser Doppler monitoring alone and in combination with MR angiography (MRA) and diffusion weighted images (DWI). Twenty-eight animals presented a successful occlusion and reperfusion determined by Doppler monitoring with an infarct size at 24 hours of 16.71±11.58%. However, when artery occlusion and infarct damage were analyzed in these animals by MRA and DWI, 15 animals were excluded and only 13 animals were included based on Doppler and MRI inclusion criteria, with an infarct size of 21.65±6.15% at 24 hours. These results show that laser Doppler monitoring is needed but not enough to guarantee a reproducible infarct volume in rat ischemic model.

**Summary statement:** Laser Doppler monitoring in combination with DWI and MR angiography represents a reliable inclusion protocol during ischemic surgery for the analysis of protective drugs in the acute phase of stroke.

## INTRODUCTION

The Stroke Therapy Academic Industry Roundtable (STAIR) criteria have been updated periodically since their creation with the purpose to improve the quality of preclinical studies of acute stroke therapies (Saver et al., 2013; STAIR, 1991). One of the most critical STAIR criteria is the monitoring of the cerebral blood flow (CBF) by Doppler during surgery with the aim to guarantee a proper medial cerebral artery occlusion (MCAo) and reproducible infarct size, usually determined at 24 hours by MR images (MRI) or histology techniques (Saver et al., 2013; STAIR, 1991).

However, in those preclinical studies focused on protective strategies for acute phase (less than 12 hours), Doppler flow monitoring is the only inclusion criteria used before treatment administration to confirm a successful ischemic damage at 24 hours, and sometimes researchers cannot really be sure if the infarct reduction might be due to the drug effect, or the variability of the model. In fact, many researchers demonstrate the efficacy of the protective drugs based on that all animals included in the study before treatment administration had a reduction of cerebral blood flow higher than 70/80% respect to the basal levels during MCAo. However, it is well known that, after MCAo, cerebral collateral circulation can supply blood to the ischemic region that is difficult to register with the Doppler probe and increases the internal variability of the experimental groups (Cuccione et al., 2016).

In this study we pretend to analyze for the first time the use of laser Doppler monitoring alone and in combination with diffusion-weighted image (DWI) and MR angiography (MRA) during MCAo and determine the infarct size variability at 24 hours in both protocols.

## RESULTS

In the surgery protocol without pterygopalatine artery ligation, a total of 34 of animals were included (Fig. 1). Initially, 6 animals were excluded due to bleeding and spontaneous death during surgery. Based on the Doppler monitoring, the rest of the animals (n=28) had a successful occlusion of the MCA (<70% respect to the basal) and reperfusion (60 min after occlusion). However, when these 28 animals were analyzed by MRA during artery occlusion, 5 animals were excluded because they had occluded the MCA and the anterior cerebral arterial (ACA) (Fig. 2), and when DWI was performed in the 23 animals selected, 10 animals were discarded because the infarcted regions were out of the established range (25-45%) (Fig. 3). DWI volume of these 13 final animals included was 33.72±6.62%. Analysis of the ischemic damage determined at 24 hours showed that the infarct size in those animals included following only laser Doppler criteria was 16.71±11.58%, while in those animals included based on Doppler criteria in combination with MRI analysis, the infarct size was significantly higher and with lower variability; 21.65±6.15% (p<0.05).

**Figure 1.**
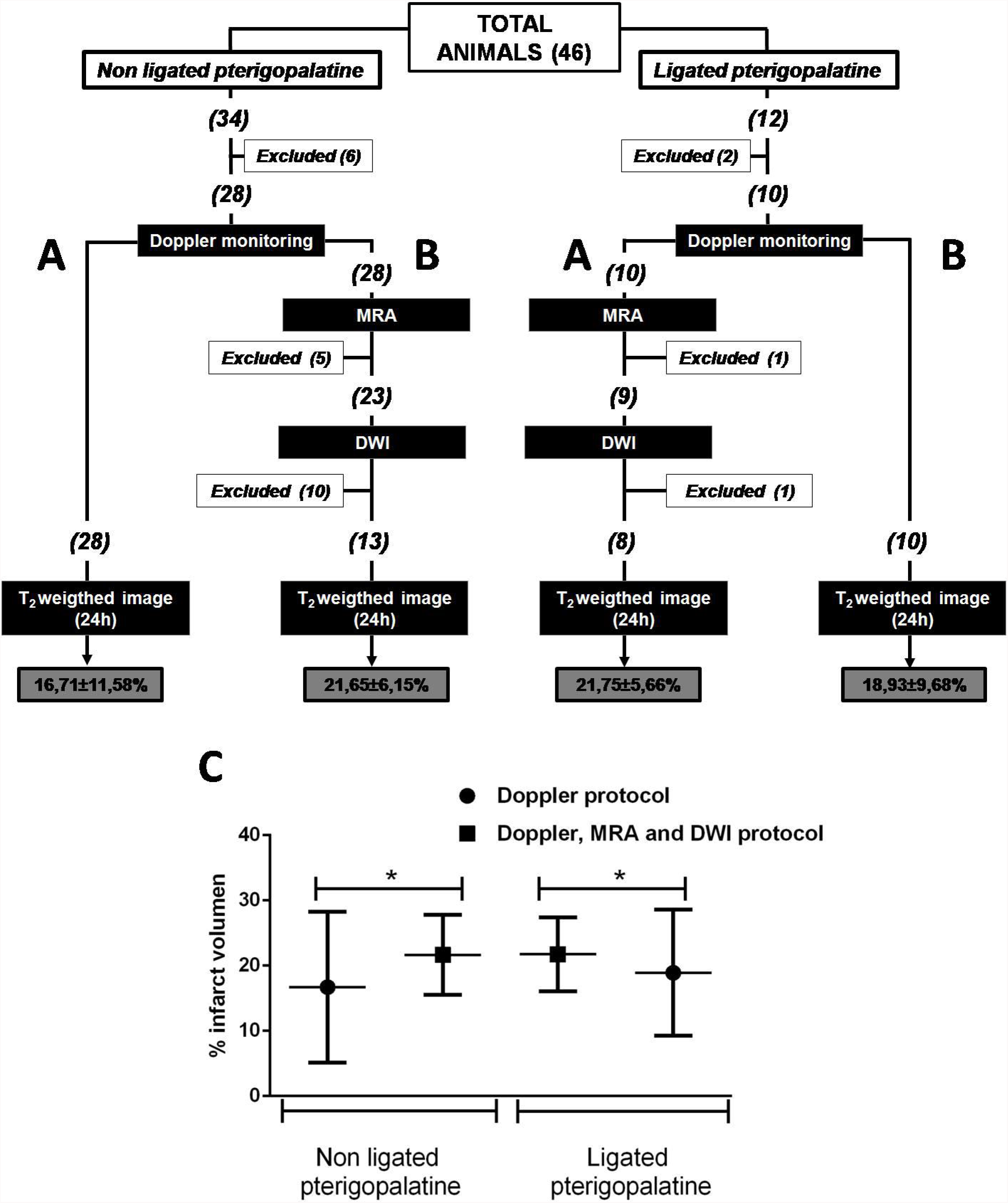
Protocol diagram summarizing the number of animals included, with exclusion per group, for final analysis. Two different experimental groups were compared: A) animals with only laser Doppler monitoring and B) animals with laser Doppler monitoring, MRA and DWI. Both groups were compared between animals with pterygopalatine with and without occlusion. C) Comparative analysis of infarct volume determined at 24 h after ischemia between the two-inclusion protocols used in rats with and without pterygopalatine occluded. Data are presented as the mean and standard error (mean ± SEM). *p<0.05.

**Figure 2.**
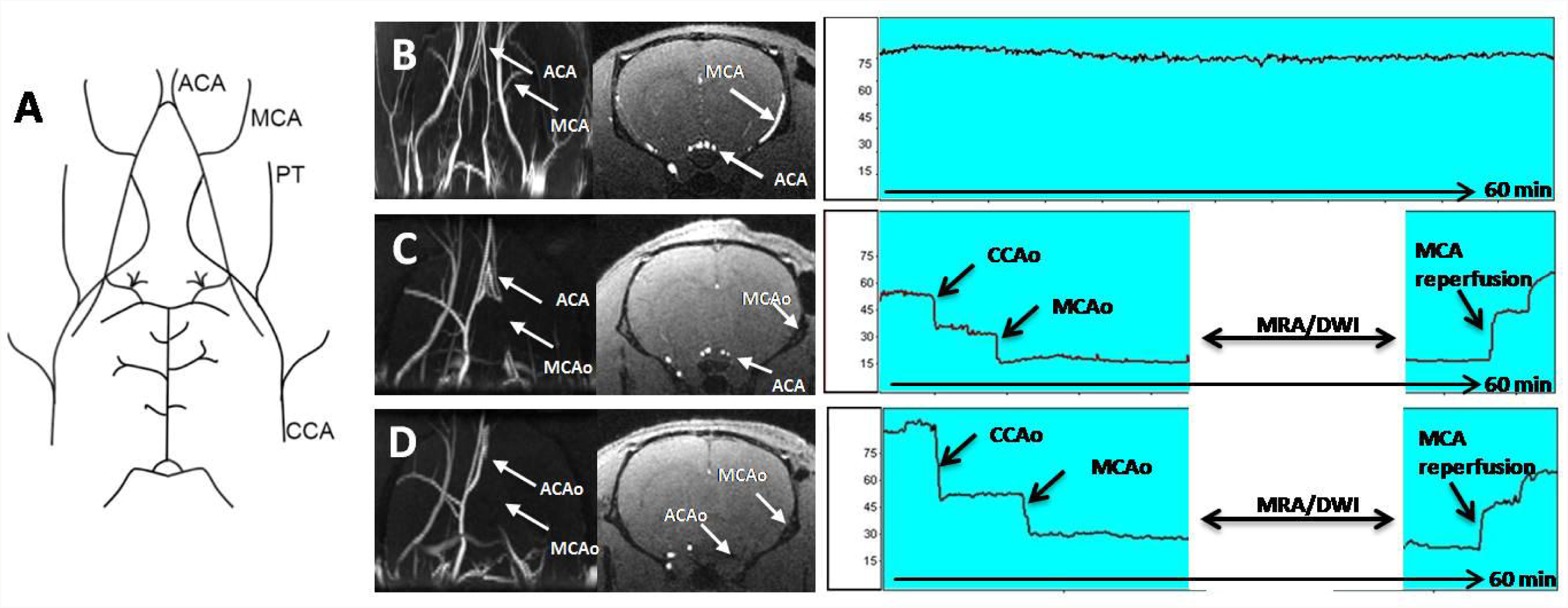
Scheme and images explaining the main aspects concerning to angiography technique. A) Diagram of cerebrovascular anatomy of the rat. Abbreviations: anterior cerebral artery (ACA), anterior cerebral artery occlusion (ACAo), middle cerebral artery (MCA), middle cerebral artery occlusion (MCAo), pterigopalatin artery (PT), common carotid artery (CCA), common carotid artery occlusion (CCAo). B) Coronal projection of a MR angiography of a healthy rat. ACA and MCA can be observed in the MR angiography projection and in the axial image below. C) Ischemic animal with the MCA and the ACA occluded, and. D) Ischemic animal with only the MCA occluded. In the laser Doppler recording is showed like animals with MCA occluded and MCA and ACA occluded showed same CBF monitoring profile during artery collusion and reperfusion.

**Figure 3.**
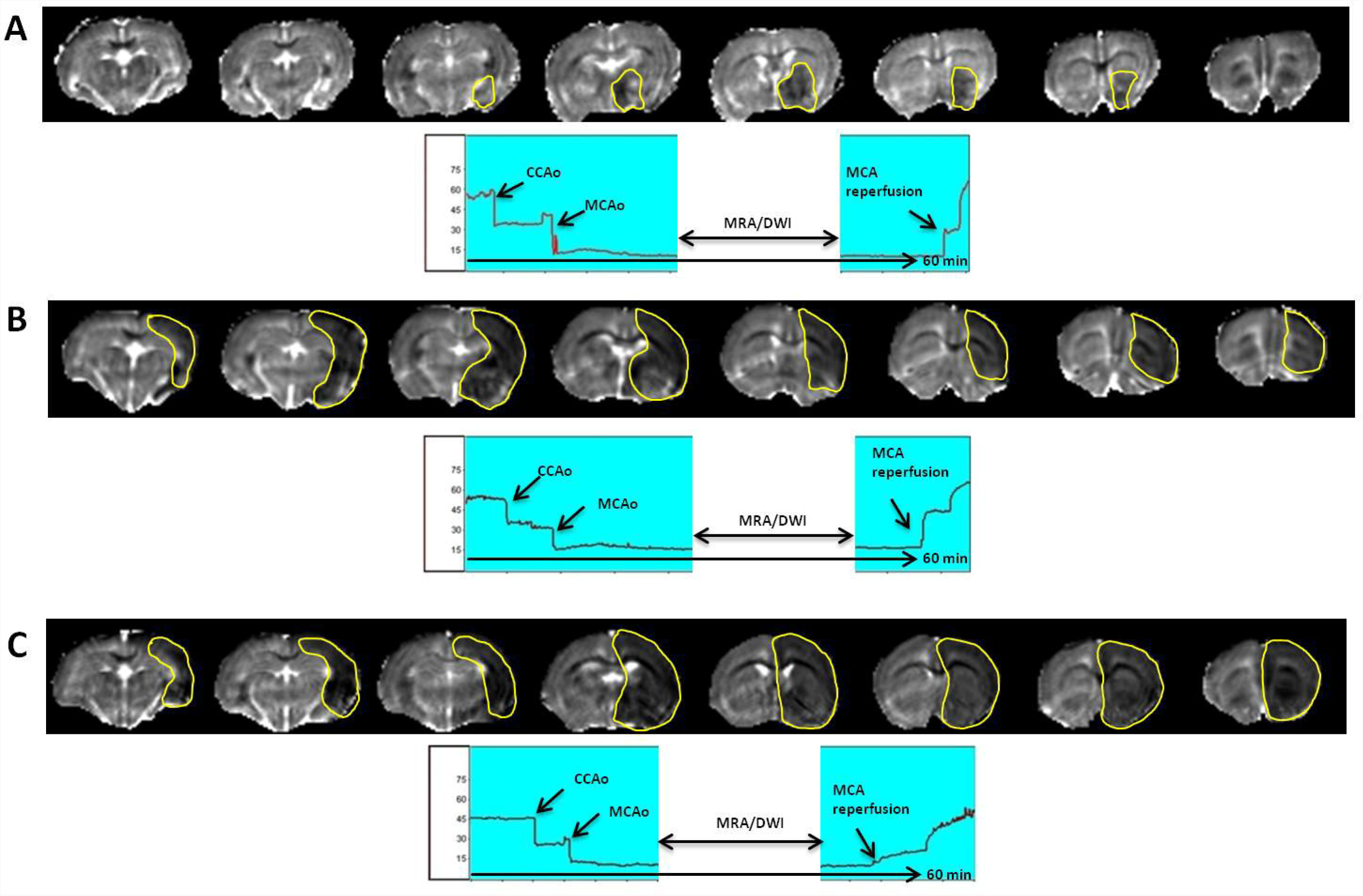
Representative ADC maps (obtained from DWI) of animals included or excluded in the study. **A)** ADC maps of an animal excluded owing to a baseline lesion volume smaller than 25% of the ipsilateral hemisphere. **B)** ADC maps of an included animal with a baseline lesion volume within the accepted range (25–45% of the ipsilateral hemisphere). **C)** Representative ADC maps of an animal excluded owing to a baseline lesion volume greater than 45% of the ipsilateral hemisphere. In the laser Doppler recording is showed like animals with different DWI volume present same CBF monitoring profile during artery collusion and reperfusion.

To validate if the pterygopalatine artery occlusion during ischemic surgery could be affecting the infarct variability previously observed, same procedure was performed. A total 12 animals were initially included (Fig. 1). Two animals were excluded due to complications during the surgery and the other 10 passed the Doppler criteria. When the MRI protocol was performed on these animals, one animal was discarded in the angiography analysis because the MCA and ACA were occluded and other animal because the infarct size was higher than the DWI threshold established. The average of the DWI volume in the animals included was 34.76±5.43%. The infarct size determined 24 hours later in those animals which passed the Doppler criteria was 18.93±9.68%, while this data was significantly higher and with lower variability in those animals subjected to both Doppler and MRI inclusion criteria (21.75±5.66%, p<0.05).

Those animals excluded based only on Doppler criteria did not showed ischemic lesion at 24 hours after MCAo.

## DISCUSSION

In line with a recent publication about the limitations of the translational stroke research (Dirnagl, 2016), an update of the inclusion criteria in ischemic animal models here described represents a demanded need to guarantee the efficacy of new protective drugs for the acute phase of stroke.

This protocol, based on the combination of laser Doppler monitoring with MRI demonstrates that the use of laser Doppler are needed to confirm that MCAo during surgery, but does not guarantee a reproducible infarct size at 24 hours, that is critical for drugs evaluation. As we could observe in this study, animals with identical arterial occlusion rate determined by laser Doppler and similar recovery flow after reperfusion, showed clear differences on infarct size determined through DWI and angiography analysis. Laser Doppler is a useful tool for ischemic model and highly sensitive when the laser probe is located on the MCA. However laser Doppler is not too much sensitive for measuring the collateral circulation (Cuccione et al., 2016), and therefore is not possible to exclude animals with extreme infarct size as DWI does. In addition, laser Doppler monitoring allows detecting the occlusion of the MCA when the filament gets the circle of Willis. However, once the MCA is occluded, with this technique is almost impossible to register if we are altering the ACA circulation, which is a critical issue because the ischemic damage will involve both MCA and the ACA territory. In this regard, angiography image allows excluding those ischemic animals in which the MCA and the ACA are occluded, reducing significantly the variability of the infarct size at 24 hours.

The pterygopalatine artery (PPA) is a maxillary artery that supplies deep structures of the face and, in some surgical protocols for ischemia induction, this artery is occluded to avoid accidental intubation of this vessel with the intraluminal suture. However, this procedure can significantly influence the MCAo model and cannot be ignored (Cuccione et al., 2016). Therefore this variable was also included in the experimental groups. In our analysis, we could observe that the PPA blocking during surgery is not relevant for this model, at least in terms of infarct variability.

Like all protocols, the use of MRI in combination with laser Doppler monitoring for animal inclusion criteria in experimental studies has limitations and advantages that should be cited.

Because the animal has to be moved after MCAo immediately to the MR (see surgery methods), one of the most important limitations for implementing this protocol is that the MRI facility must be close to the surgery bench to keep the animals under anesthetized conditions and reduce as much as possible the movement of the filament located in the artery; that is not very common in many research centers, and also worst when the MRI facility is in a different building respect to the surgery facilities. Other important consideration is that with Doppler monitoring is possible to achieve a rate of inclusion of 100% but with this new protocol, the inclusion rate is under 50%, which means a loss of animals, time and money. We have also established an arbitrary inclusion interval between 25 and 45% for DWI hemispheric infarct volume that could be discussed. We have established this threshold because, based on previous studies (Argibay et al., 2017; Campos et al., 2011; Pérez-Mato et al., 2014; Vieites-Prado et al., 2016) where same inclusion protocol was used, we observed that DWI volumes under 25% during MCAo were associated with small subcortical ischemia or no ischemia at 24 hours, while DWI volumes above 45% were associated with malignant infarct that affected all hemisphere and caused a high rate of mortality. Therefore, based on this experience, we established this threshold for DWI. Finally, this study was performed in Sprague-Dawley rats because this is the most common strain used in the field of cerebral ischemia, but we are aware that this protocol should be validate in other strains, in mice or also in a permanent MCAo model.

In despite of these limitations, we would like to highlight that this new inclusion protocol allows reducing significantly the infarct size variability at 24 hours after ischemia that is critical for treatment evaluation. In addition, this protocol also permits to determine a basal ischemic lesion in the animals included before treatment administration that, in combination with the ischemic size determined after treatment, enhances the quality and the reliability of the results.

In brief, we can conclude that laser Doppler monitoring is needed but not enough to guarantee a reproducible infarct volume in rat ischemic model. Laser Doppler monitoring in combination with DWI and MR angiography represents a reliable inclusion protocol during ischemic surgery for the analysis of new protective drugs focused on the acute phase of stroke.

## MATHERIAL AND METHODS

### Animals

All experimental protocols were approved by the local Animal Care Committee according to the guidelines established by the European Union (86/609/CEE, 2003/65/CE,and 2010/63/EU) and following the ARRIVE Guidelines for animal experiments. Male Sprague-Dawley rats weighing between 280 and 330 g were used (11-12 weeks old). Animals were housed individually at an environmental temperature of 23°C, with 40% relative humidity and a 12 h light-dark cycle, and were given free access to food and water.

### Rat model of cerebral ischemia and MR imaging

All surgical procedures were performed under sevoflurane anaesthesia (6% induction and 4% maintenance in a mixture of 70% NO_2_ and 30% O_2_). Rectal temperature was maintained at 37 ± 0.5°C in all animals during surgery using a thermostat-controlled electric pad (Neos Biotec, Pamplona, Spain). The animal head was placed on a porexpan plate to avoid direct contact between the pad and head. Transient focal ischemia (60 min) was induced by intraluminal occlusion of the MCA, following the method described previously (Howels et al., 2016; Lee et al., 2014). All surgeries were performed by one experienced researcher (>2 years) in the transient intraluminal filament MCAo model.

Occlusion was performed using commercially available sutures with silicone-rubber-coated heads (350p μ m in diameter and 1.5 mm long; Doccol, Sharon, MA, USA). CBF was monitored with a Periflux 5000 laser Doppler perfusion monitor (Perimed AB, J¨rf¨lla, Sweden) by placing the Doppler probe (model 411; Perimed AB) under the temporal muscle at the parietal bone surface, near the sagittal crest. Twenty-five minutes after artery occlusion had been achieved, as indicated by Doppler signal reduction (CBF reduction higher than 70%), each animal was carefully and immediately (in less than 1 min) moved from the surgical bench to the MR system for ischemic lesion assessment using diffusion-weighted images (DWI). In combination with DWI, MRA was performed to ensure that the artery remained occluded throughout the MR procedure and to confirm the only occlusion of the MCA. Animals were then returned to the surgical bench and the Doppler probe was repositioned. Reperfusion was performed 60 min after occlusion onset. Ischemic damage was confirmed and determined 24 h after ischemia by MR form T2 weighted image. The transient intraluminal filament MCAo model was used because represents the most common model used in stroke experiments field (Cuccione et al., 2016).

### Experimental groups

Two different experimental inclusion criteria were compared.

- Animals with CBF reduction higher than 70% and complete reperfusion after MCA occlusion determined only by laser Doppler monitoring.
- Animals with CBF reduction higher than 70% determined by laser Doppler monitoring, DWI hemispheric infarct volume between 25 and 45 % (indicated as % of ischemic damage respect to the ipsilateral hemisphere volume), MRA of the occlusion of the MCA, and complete reperfusion after MCA occlusion.

Both inclusion protocols were also compared with the PPA occluded.

### MRI Imaging

All studies were conducted on a 9.4T horizontal bore magnet (Bruker BioSpin, Ettlingen, Germany) with 440 mT m^−1^ gradients and a combination of a linear birdcage resonator (70 mm in diameter) for signal transmission and a 2×2 surface coil array for signal detection. MRI post-processing was performed using ImageJ software (W. Rasband, NIH, USA).

Basal ischemic lesion during MCA occlusion was determined by counting pixels with apparent diffusion coefficient (ADC) values below a threshold in the ipsilateral brain hemisphere. The values of ADC in the healthy rat brain normally do not fall below 0.55 × 10^−3^ mm^2^ s^−1^; therefore, this threshold provides a convenient means of segmenting abnormal tissue (Reith et al., 1995). ADC maps were obtained from diffusion-weighted images (DWI) using a spin echo echo-planar imaging sequence (DTI-EPI) with the following acquisition parameters: echo time (TE) = 26.91 ms, repetition time (TR) = 4 s, spectral bandwidth (SW) = 200 KHz, 7 b-values of 0, 300, 600, 900, 1200, 1600, and 2000 s/mm^2^, flip angle (FA) = 90°, number of averages (NA) = 4, 14 consecutive slices of 1 mm, field of view (FOV) = 24 × 16 mm^2^ (with saturation bands to suppress signal outside this FOV), a matrix size of 96 × 64 (isotropic in-plane resolution of 250 μm/pixel × 250 μ m/pixel) and implemented with fat suppression option.

To evaluate the status of MCA occlusion in a noninvasive manner, the time-of-flight magnetic resonance angiography (TOF-MRA) was performed. The TOF-MRA scan was performed with a 3D-Flash sequence with an echo time (TE) = 2.5 ms, repetition time (TR) = 15 ms, flip angle (FA) = 20°, number of averages (NA)= 2, spectral bandwidth (SW)= 98 KHz, 1 slice of 14 mm, field of view (FOV) = 30.72×30.72×14 mm^3^ (with saturation bands to suppress signal outside this FOV), a matrix size of 256×256× 58 (resolution of 120 μm/pixel×120 μ m/pixel×241 μ m/pixel) and implemented without fat suppression option. DWIs and TOF-MRA were acquired during MCAo, at the same time (30±5 min after occlusion).

Ischemic lesions were determined at 24 h after ischemia from T2 maps. These maps were calculated from T2 weighted images using a multi-slice multi-echo sequence (MSME) with echo time (TE)= 9 ms, repetition time (TR)= 3 s, 16 echoes with 9 ms echo spacing, flip angle (FA) = 180°, number of averages (NA)= 2, spectral bandwidth (SW)= 75 KHz, 14 slices of 1 mm, field of view (FOV) = 19.2 × 19.2 mm^2^ (with saturation bands to suppress signal outside this FOV), a matrix size of 192× 192 (isotropic in-plane resolution of 100 pm/pixel × 100 μ m/pixel) and implemented without fat suppression option. Infarct size was indicated as % of ischemic damage respect to the ipsilateral hemisphere volume, corrected for brain edema. The image evaluations were performed by a blind researcher.

### Statistical analysis

All data are presented as the mean and standard error (mean ± SEM). The Student-test was used to identify significant differences between two groups. Statistical significance was set at P<0.05. Statistical analyses were conducted using SPSS Statistics for Macintosh, Version 18.0 (IBM, Armonk, NY, USA).The statistical analysis was performed by a blind researcher.

## COMPETING INTEREST

No competing interest declared

## AUTHOR CONTRIBUTIONS

José Castillo and Francisco Campos designed research. Héctor Fernández, Ramón Iglesias, Antonio Dopico performed the experiments. Maria Pérez and Tomás Sobrino analysed data, and José Castillo, Héctor Fernández and Francisco Campos wrote the manuscript.

## FUNDING

This study has been partially supported by grants from Instituto de Salud Carlos III (PI13/00292; PI14/01879). Furthermore, T. Sobrino (CP12/03121) and F. Campos (CP14/00154) are recipients of a research contract from Miguel Servet Program of Instituto de Salud Carlos III. The funders had no role in the study design, data collection and analysis, decision to publish, or preparation of the manuscript.

